# Network level enrichment provides a framework for biological interpretation of machine learning results

**DOI:** 10.1101/2023.10.14.562358

**Authors:** Jiaqi Li, Ari Segel, Xinyang Feng, Jiaxin Cindy Tu, Andy Eck, Kelsey King, Babatunde Adeyemo, Nicole R. Karcher, Likai Chen, Adam T. Eggebrecht, Muriah D. Wheelock

**Affiliations:** Department of Mathematics and Statistics, Washington University in St. Louis, MO 63130, USA; Department of Radiology, Washington University in St. Louis, MO 63110, USA; Department of Neurology, Washington University in St. Louis, MO 63110, USA; Department of Psychiatry, Washington University in St. Louis, MO 63110, USA

**Keywords:** functional connectivity, machine learning, twins, feature selection, brain networks, HCP

## Abstract

Machine learning algorithms are increasingly used to identify brain connectivity biomarkers linked to behavior and clinical outcomes. However, non-standard methodological choices in neuroimaging datasets, especially those with families or twins, have prevented robust machine learning applications. Additionally, prioritizing prediction accuracy over biological interpretability has made it challenging to understand the biological processes behind psychopathology. In this study, we employed a linear support vector regression model to study the relationship between resting-state functional connectivity networks and chronological age using data from the Human Connectome Project. We examined the effect of shared variance from twins and siblings by using cross-validation, either randomly assigning or keeping family members together. We also compared models with and without a Pearson feature filter and utilized a network enrichment approach to identify predictive brain networks. Results indicated that not accounting for shared family variance inflated prediction performance, and the Pearson filter reduced accuracy and reliability. Enhancing biological interpretability was achieved by inverting the machine learning model and applying network-level enrichment on the connectome, while directly using regression coefficients as feature weights led to misleading interpretations. Our findings offer crucial insights for applying machine learning to neuroimaging data, emphasizing the value of network enrichment for comprehensible biological interpretation.

## 1. Introduction

Supervised machine learning (ML) is applied increasingly to understand how human brain connectivity supports behavior, cognition, and emotion, as well as how aberrant brain connectivity leads to psychopathology (Cui & Gong, 2018; Hahn et al., 2022; Nielsen et al., 2019). Among modern ML techniques, linear regression models have found most success in human neuroimaging research, e.g. Ridge regression and Linear Support Vector Regression (LSVR) with a Ridge penalty (Cui & Gong, 2018; Dhamala et al., 2020; Taxali et al., 2021; Vergun et al., 2013). However, several challenges remain for practical application and interpretation of ML models in neuroscience research. First, a considerable number of neuroscience studies are designed to leverage twins and families with heritable characteristics, yet there is a lack of guidance or standardized approach to modeling shared variance among related individuals (Cohen et al. 2020; Elliott et al. 2019; Feilong et al. 2021). Further, ML studies frequently report prediction accuracy, but fail to evaluate the biological validity or fundamental brain mechanisms leading to clinical and behavioral outcomes (He et al. 2020; Niu et al. 2020; Wang et al. 2013). Finally, while feature selection procedures may increase computational efficiency, the impact on reliability and biological interpretability of ML models are poorly understood (Dadi et al., 2019; Nielsen et al., 2019; Shi et al., 2021).

Resting-state functional magnetic resonance imaging (rs-fMRI) is a non-invasive method of assessing brain connectivity (Zhang & Raichle, 2010) via identifying synchronous blood oxygen-level dependent (BOLD) changes in multiple brain regions while subjects lie in the scanner but do not perform a task. Resting-state functional connectivity (rsFC) can be derived from the temporal correlation of spontaneous fluctuations of such BOLD signals between brain regions (Biswal et al., 1995) and estimating rsFC across all possible regions in the brain forms the basis of the functional connectome (Ombao et al., 2016). Notably, brain regions that are correlated have been reproducibly classified into resting-state networks (Gordon et al., 2016; Power et al., 2012; Yeo et al., 2011). Previous studies have shown that these functional networks in the brain are disrupted in patients with neurological disorders (e.g., Alzheimer disease, Parkinson’s, multiple sclerosis, stroke, etc.) (Wheelock et al. 2023) and psychiatric disorders (e.g., anxiety, internalizing, externalizing, neuroticism, etc.) (Menon, 2011). Understanding the disruption of these networks is therefore of putative clinical importance. ML statistical techniques offer great promise for predicting complex psychiatric and neurologic dysfunction using rs-fMRI data (Arbabshirani et al., 2017; Bhaumik et al., 2017; Cropley et al., 2021; Guo et al., 2012). However, several methodological gaps have posed challenges when effectively implementing and interpreting ML models in human neuroimaging data.

Shared variance due to families or longitudinal designs (repeated measures within a subject) poses challenges to ML cross-validation modeling. Related subjects like family members are often included in human rs-fMRI neuroimaging datasets. For example, twins and siblings are present in developmental cohorts such as the Infant Brain Imaging Study (Eggebrecht et al., 2017), Adolescent Brain Cognitive Development (ABCD) study (Hahn et al., 2022), and the Human Connectome Project (HCP) (Van Essen et al. 2012; Cohen et al. 2020; Greene et al. 2020; Feilong et al. 2021). However, this shared variance is rarely properly dealt with in ML studies (Cohen et al., 2020; Cui & Gong, 2018; Scheinost et al., 2019). Several attempts have been made to employ random sampling with cross-validation to avoid the issue of shared variance among related individuals (Winkler et al., 2015). One such approach is to reduce the total sample size by only including non-related individuals (Dhamala et al. 2020; Tian and Zalesky 2021; Gbadeyan et al. 2022). For instance, some researchers retain only one member from each family (Li et al. 2017; Nostro et al. 2018; Elliott et al. 2019; Seitzman et al. 2020; Lohmann et al. 2021). Nevertheless, this type of method drastically reduces prediction power due to a decreased number of subjects, which might defeat the purpose of twin study designs. Alternatively, studies carefully considering shared variance in the cross-validation step have implemented a leave-one-family-out cross-validation scheme in which related individuals are only in either the training or testing set but not both (Dubois et al. 2018; He et al. 2020; Feilong et al. 2021). Though a few solutions to shared variance among family members have been proposed, a lack of consistent standards for modeling such shared variance remains a hindrance to the reliable and well powered study of neuroimaging datasets.

A second challenge arises from the vast number of features typically found in rsFC across the entire brain, amounting to several thousand, necessitating the need to address issues such as overfitting and excessive computational requirements in many ML models. Consequently, an additional feature selection step is frequently implemented prior to the ML model in order to reduce the number of FC features included in the regression model (Arbabshirani et al., 2017; Craddock et al., 2009; Gao et al., 2019). Many feature selection methods fall into the category of massively univariate approaches which model the relationship between each FC feature and clinical outcome independently, such as marginal Pearson screening with the top-ranked features selected for inclusion in the ML model (Nielsen et al. 2019; Fan et al. 2006; Shen et al. 2017). However, a notable concern of such feature selection methods is that they may exclude features that have a significant contribution in a full multiple regression model, particularly when the predictors exhibit high correlation (Wang et al. 2020).

A third and significant challenge of ML models is that biologists, neuroscientists, and clinicians also demand interpretable models since dysfunction within specific brain systems could differentially contribute towards a behavior or clinical outcome. Yet, biological interpretation of ML results remains a challenge for myriad reasons. While studies have interpreted the beta weights of features from ML models (Plitt et al. 2015; Finn et al. 2015; Dhamala et al. 2020; Jiang et al. 2020; Greene et al. 2020; Bellantuono et al. 2021; Kardan et al. 2022), it has been shown that such interpretation could be seriously misleading because some features that are not related to the target label can still have significant weights in prediction (Chen et al., 2022; Haufe et al., 2014). For example, features with the strongest ML beta weights may reflect non-neuronal or nuisance signal such as head motion or machine noise (Chen et al., 2022; Haufe et al., 2014; Siegel et al., 2017). In other words, beta weights from ML models only represent a combination of features for optimal prediction performance and should not be confused with the biological relation to the behavioral/clinical vector labels. One method to resolve this issue of interpretability of the rsFC edge-level model weights is to first calculate the covariance between the predicted label and the rsFC between a pair of regions of interest (ROI), hereafter referred to as the inversion model (first introduced by Haufe et al. (2014) to the neuroimaging community). A second method has leveraged the systems level architecture of the brain as described in several canonical atlases (Power 2011; Gordon 2016). These canonical atlases can be used to describe within-network and between-network connectivity - herein referred to as “network block” connectivity. One commonly employed approach is an “individual network block” method in which only functional connections within a single network block (i.e., either within or between network rsFC) are used as features within the model (Kardan et al., 2022; Millar et al., 2020, 2022; Nielsen et al., 2019; Rudolph et al., 2018). Given the fact that increasing feature count is commonly associated with increasing prediction accuracy, limiting ML models to only contain features within specific networks has resulted in limited accuracy (Bellantuono et al., 2021; Cui & Gong, 2018; Nielsen et al., 2019) and reliability (Mellema et al., 2021; Tian & Zalesky, 2021). An alternative approach, Network Level Analysis (NLA), has been recently introduced to assess the network-level enrichment of connectome-wide associations, precluding any need to limit the number of features in the ML model. NLA has been successfully applied in several univariate association papers to understand connectome-wide associations with attention (Wheelock et al. 2021), emotion (Gilbert et al., 2021; Perino et al., 2021), gestational age (Wheelock et al. 2019), and developmental behaviors associated with autism (Eggebrecht et al., 2017; Marrus et al., 2018; McKinnon et al., 2019), but has not been applied to ML models. The combined computational pipeline of NLA + ML with the inversion model properly embedded has great potential for both predicting outcomes with increasing prediction accuracy and facilitating biological interpretation.

To summarize, this paper provides benchmarks for the most effective ML methods applicable to twin and family study designs, facilitating a consistent guideline for ML modeling using rsFC data with enhanced prediction accuracy, reliability and biological interpretability. First, we tested the LSVR model with and without a feature filter and contrasted that with simple Pearson correlations between rsFC and age. Further, to account for the shared variance, we derived a cross-validation pipeline where a random-sampling scheme is utilized such that family members were not split across folds and stayed together within training or test set for each sampling round. As a comparison, we applied another cross-validation method that randomly assigned all the subjects into training and test sets. The prediction accuracy, accounted variance of the model, and reliability were evaluated as the performance measure for each model. Specifically, we fit ML models with the datasets from different days as inputs and visualized the measures for predictive performance as well as the features contributing significantly to the prediction. Finally, we evaluated two methods that leverage the systems level organization of the brain to aid in biological interpretation. Specifically, we assessed the performance of fitting ML models in an individual network block analysis and established the feasibility and utility of the novel combination of ML and NLA methods.

## 2. Materials and Methods

### 2.1 Data Characteristics

The publicly available dataset from the HCP S1200 release was considered in the present study (Glasser et al., 2013). HCP study design included recruitment of twins and family members. Behavior was assessed on one day and two 30 minute scans of resting state fMRI were acquired on two separate days (here referred to as Rest 1 and Rest 2). This study design allowed for the assessment of test-retest reliability. A total of 965 healthy adults (ages 22-35 years old) were identified as having at least 10 minutes of low motion data for both Rest 1 and Rest 2 of the HCP rs-fMRI data and were included for further analysis. Among all 965 subjects, there were 420 families with a maximum of 5 members per family.

### 2.2 Data Acquisition

High-resolution T1-weighted (MP-RAGE, 2.4s TR, 0.7×0.7×0.7mm voxels) and BOLD contrast-sensitive (gradient echo EPI, multiband factor 8, 0.72s TR, 2×2×2mm voxels) images were acquired from each participant using a custom Siemens SKYRA 3.0T MRI scanner and a custom 32 channel Head Matrix Coil. The HCP employed sequences with both left-to-right (LR) and right-to-left (RL) phase encoding, with each participant completing a single run in each direction on two consecutive days, which results in a total of four runs including two runs for Rest 1 and another two for Rest 2 (Van Essen et al., 2012).

### 2.3 Data Processing

Minimally pre-processed data have been shown to be insufficient in controlling for confounds such as subject head motion (Burgess et al. 2016). Additional research suggests that sufficient low motion functional connectivity data must be available for each subject in order to make reliable claims about associations between functional connectivity and behavior or outcomes (Gordon, Laumann, Gilmore, et al., 2017; Laumann et al., 2015). In this section, we provide the preprocessing steps of the dataset.

#### 2.3.1 FC Preprocessing

The HCP functional MRI data preprocessing methods employed in this study have been previously described (Gordon et al. 2017). First, to account for magnetization equilibrium and any responses evoked by the scan start (Laumann et al., 2015), the first 29.52 seconds or -- 41 frames -- of each resting-state run were discarded. Then, the functional data were aligned to the first frame of the first run using rigid body transforms, motion corrected (3D-cross realigned), and whole-brain mode 1000 normalized (Miezin et al., 2000). The data, consisting of 2×2×2mm voxels, was then registered to the T1-weighted image and a WashU MNI atlas using affine and FSL transforms (Smith et al., 2004).

Further preprocessing of the resting-state BOLD data was applied to remove artifacts (Ciric et al., 2017; Power et al., 2014). Specifically, frame-wise displacement (FD), a metric used to quantify the amount of motion or displacement between consecutive frames in fMRI data, was calculated (Power et al., 2012) and artifact removal (Ciric et al., 2017; Power et al., 2014) was completed with a low-pass filter at 0.1 Hz to address respiration artifacts affecting the FD estimates (Fair et al., 2020; Siegel et al., 2017), along with a threshold after the low-pass respiration filter to remove frames with FD greater than 0.04 mm (Dworetsky et al., 2021). To prepare the data for functional connectivity (FC) analysis, the regression of nuisance variables was performed, including: 1) whole-brain mean, 2) ventricular and white matter CSF signals, 3) the temporal derivatives of those regressors, and 4) 24 movement regressors (Friston et al., 1996; Satterthwaite et al., 2012; Yan et al., 2013). Temporal masks of the gray matter, white matter, and ventricles were created from the T1-weighted images for each of the individual-specific regressors using Freesurfer 5.3 automatic segmentation (Fischl et al., 2002). Segments of data lasting fewer than 5 contiguous frames were excluded, and then least squares spectral estimation was used to interpolate over the censored frames (Hocke & Kämpfer, 2008; Power et al., 2014). Data were then bandpass filtered from 0.009 to 0.08 Hz, and censored frames were removed from the time series (Seitzman et al., 2020).

Following previously established methods (Gordon et al., 2016), the preprocessed BOLD time series data underwent surface processing, which involved using the ribbon-constrained sampling procedure in Connectome Workbench to sample the BOLD volumes to each subject’s individual native surface and exclude voxels with a time series coefficient with a variation 0.5 SDs above that of the mean of nearby voxels (Glasser et al., 2013; Gordon et al., 2016). After being sampled to the surface, time courses were then deformed, resampled, and smoothed using a Gaussian smoothing kernel (FWHM = 4mm, sigma = 1.7). Connectome Workbench was then used to combine these surfaces with volumetric subcortical and cerebellar data into the CIFTI format to create full brain time courses, excluding non-gray matter tissue (Glasser et al., 2013).

#### 2.3.2 Whole-Brain rsFC Feature Extraction

After the processing procedure above, surface-based parcels and canonical functional networks (Gordon et al., 2016) were used to parcellate a set of 333 previously defined regions of interest (ROIs) into 12 networks and one unspecified network as shown in Figure 1A, where the unspecified network consists of unassigned parcels that were not strongly connected with any other parcels as defined in Gordon et al. 2016. For each subject, the mean time series within each ROI was calculated by taking the average of time series over all the voxels within this ROI. Since we randomly sampled 10 minutes of low-motion frames from each ROI for each subject on two scan days, all 965 subjects had the same amount of low-motion data on both days. The Pearson correlation between the mean time courses of each pair of ROIs was evaluated (55,278 pairs in total, ROIs on the diagonal excluded) and normalized with a Fisher-z transformation. A correlation matrix consisting of these normalized correlations was constructed for each of the two scan days, respectively, across all the 965 participants. For each scan day, the average and standard deviation of all 965 correlation matrices are shown in Figure 1B.

**Figure 1.**
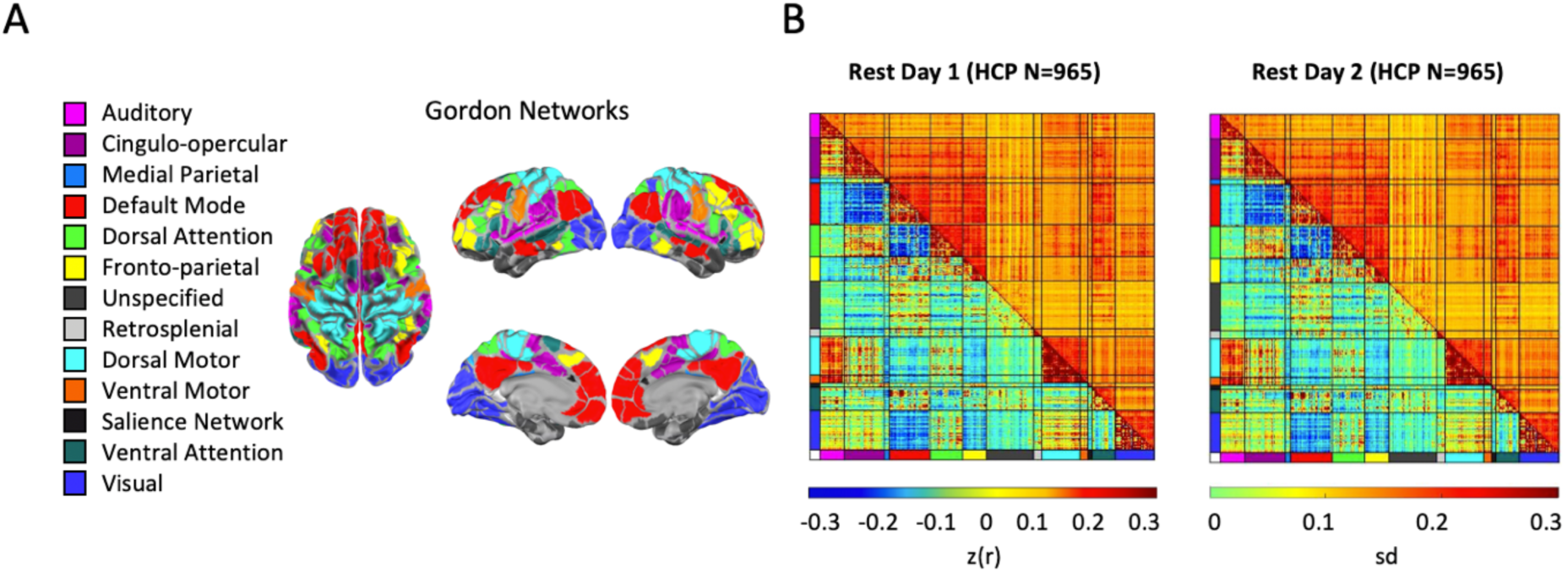
Resting-state functional connectivity (rsFC) data. **(A)** Gordon networks. As defined in Gordon et al. 2016, 333 parcels in the brain were used to extract mean rsFC and were grouped into 12 networks and one unspecified network. **(B)** Mean and standard deviation for rsFC on two separate days. For each pair of 333 parcels, the mean (lower triangle) and the standard deviation (upper triangle) for the 10 minutes of low-motion data rsFC data were computed (normalized to z-scores) across all 965 HCP subjects on two different scan days.

### 2.4 Machine Learning Model

#### 2.4.1 Linear Support Vector Regression (LSVR)

Rather than fitting a linear regression model on each feature independently, we instead utilized a linear support vector regression (LSVR) to model the relationship between multiple features (i.e., an individual’s rsFC) and chronological age. Note that support vector regression (SVR) is an extension to support vector machine (SVM), which is frequently applied to classification problems with binary labels, such as testing if an individual can be identified as a part of a specific category based on the rsFC. Since the predicted response in our case (i.e., chronological age) is a continuous variable, we employed SVR to fit an ML regression model and predict the continuous label. Various kernel functions can be chosen when applying SVR, such as linear kernels, sigmoid kernels and Gaussian kernels (RBF). Among them, Linear SVR is commonly employed in rsFC studies since it is less prone to overfitting than non-linear ones and is significantly faster to train (Cui and Gong 2018; Taxali et al. 2021; Wang et al. 2013).

For all approaches detailed below, we divided the data using an 80% training and 20% test hold out procedure and repeated this 1000 times. The hyper-parameter in the LSVR model (i.e., the tuning parameter of the L^2^ Ridge penalty term) was selected by a 5-fold cross-validation within the training data on each iteration. Specifically, on each iteration within the training set containing 80% of the subjects, we further randomly divided the data into an inner-cross-validation training set with 80% subjects and an inner-cross-validation test set with 20% subjects. We tried different hyper-parameters in a certain range to fit the LSVR model using the inner-training set and picked the optimal one with the minimized mean square error (MSE) yielded by the prediction using inner-test set for each iteration. We refer to Figure S1 in the supplementary materials for the clarification of this nested 5-fold cross-validation method. To evaluate the performance of LSVR models, we adopted three different measures: 1) prediction accuracy (r), which is the Pearson correlation between the predicted and actual ages, 2) explained variance (R^2^), and 3) reliability score of brain-behavior predictions, that is the intra-class correlation statistic (ICC). Specifically, we choose the type ICC(2,1) in this study because we are predicting a behavior label, and our goal is to evaluate the reliability of brain-behavior predictions instead of the reliability of rsFC (Koo & Li, 2016; Taxali et al., 2021). We also reported the MSE and the mean absolute error (MAE) in the supplementary materials as two additional measures of prediction accuracy. For each iteration, we first fit the ML models using the training set and then calculated the evaluation measures above in the test set by comparing the predicted labels and the actual ones.

#### 2.4.2 Random Sampling and Cross-Validation Scheme

To test the impact of shared variance across family members on the ML model performance, we employed two different random sampling approaches. In the first sampling approach, we assigned participants to the training or test set randomly ignoring family structure, an approach that is commonly used across datasets including HCP (Cohen et al., 2020; Cui & Gong, 2018; Rasero et al., 2018). In the second sampling approach, we used an identical 80-20% training-test split with 1000 folds but assigned families to either the training or test set but not both (i.e., family members were never in both the training and testing sets) in order to account for the family structure within the HCP dataset (Figure 2).

**Figure 2.**
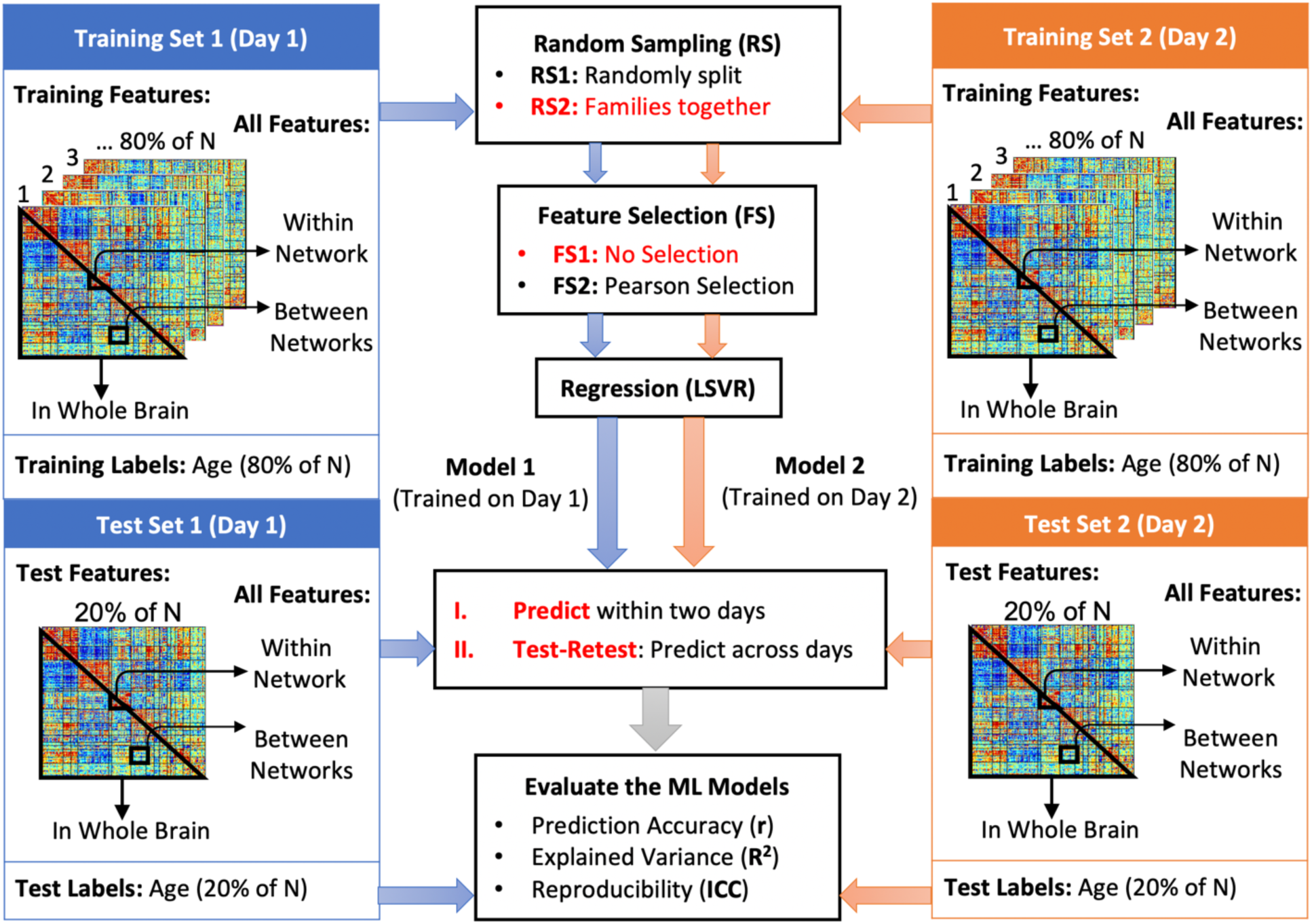
Overview of the Machine Learning (ML) pipeline. Linear support vector regression (LSVR) was employed to fit a regression model for the relationship between rsFC matrices and the age labels on two separate scan days (Rest day 1 and Rest day 2). On each day, we randomly sampled 80% of the subjects, assigning them to a training set, and the remaining 20% subjects were assigned to a test set. Two different random sampling (RS) mechanisms were applied for comparison. One is to randomly split the subjects to the training and test sets without considering the family structure (RS1), and another one is to keep family members together within either training or test set (RS2). Both random sampling procedures were repeated for 1000 iterations, respectively. Within the training set of each iteration, the hyper-parameter was chosen by an embedded 5-fold cross-validation. We refer to Figure S1 in the Supplementary materials for the details of this nested cross-validation (CV). For the test-retest, we trained an LSVR model on Rest 1, predicted the ages on Rest 2, and fit another LSVR model via the training set from Rest 2 and tested on Rest 1, respectively. We averaged the prediction accuracy of two predicted sets to yield the ICC(2,1).

#### 2.4.3 Feature Selection

In order to understand the impact of feature selection on prediction accuracy, reliability, and biological interpretability, we employed two feature filter approaches (Figure 2). In the first approach, we employed a feature filter on the training set in which we ranked features according to the marginal Pearson correlation between each functional connection and the age label for all subjects in the training set. Then, the 1,000 top ranking features (strongest univariate correlations between rsFC and age) were selected to enter the LSVR model since the performance of an LSVR model would converge when the number of predictors surpassed 1,000 (Cui & Gong, 2018; Nielsen et al., 2019). We continued to fit the LSVR model by the same training set that we used for feature selection to ensure that, in each 1000 folds, the test data was never touched before prediction. In the second approach, we applied no feature filter and allowed all edges within the lower triangle of the rsFC matrix, excluding the diagonal, to enter into the LSVR model (55278 features).

Further, given that Pearson correlation is also frequently used as a univariate method to investigate the relationship between the connectivity matrices and the labels (i.e., age), we further compared the significant features selected by the univariate correlation approach (i.e., Pearson) and the multivariate regression model (i.e., LSVR) with and without a Pearson feature filter. Finally, as pointed out by Haufe et al. (2014), an inversion process on the ML results would be necessary for the biological interpretation and shall yield similar significant features as the Pearson feature filter. Thus, we applied the inversion model together with the LSVR model to obtain the inverted weights of each feature. To summarize, we calculate the estimated weights for each ROI pair by four different methods: 1) Pearson correlation, 2) Pearson correlation feature filter + LSVR, 3) LSVR without feature filter, and 4) LSVR with inversion. Specifically, for the first method, we calculated the Pearson correlation between each feature in rsFC and the age label. For the second method, we applied the Pearson feature filter ahead of LSVR model fitting as we described in the last paragraph. Then we trained the LSVR model and estimated the beta weights of the features that included in the model (i.e., the features that have been selected in the screening step). For the third approach, we estimated the beta weights of all 55278 features by the regression coefficients in the LSVR model. For the last approach, we computed the covariance between each rsFC and the predicted age as the estimated inverted weight for each ROI pair. We implemented all four methods via the two sampling schemes (random and accounting for families), respectively, and we reported the weights averaged over 1000 replications.

#### 2.4.4 Interpretation of ML Regression Weights

In contrast to methods such as Pearson correlation or ordinary least squares regression which fall in the category of forward models in the multivariable data analysis (Haufe et al., 2014), it can be severely misleading to provide biological interpretations for the regression weights (Jiang et al., 2020; Sripada et al., 2019) or selected features (Finn et al., 2015; Greene et al., 2018) of the backward multivariable ML models. Roughly speaking, forward models investigate how the observed variables are driven by the underlying factors, while backward models focus on expressing the variables of interest as a function of the observed data. Therefore, the interpretability of an ML model is determined by the direction of the functional relationship between observations and underlying variables: the beta weights of forward models are interpretable, while those of backward models are not in most cases. Due to this distinction, an inversion process is necessary to facilitate reasonable interpretations for the backward models. We followed the elegant inversion model (Haufe et al., 2014) to resolve this interpretational issue. Specifically, we transformed our backward LSVR model into the forward form by computing the covariance between the predicted target variable age and the rsFC (Chen et al., 2022). We point out that the rsFC-age covariance by the inversion model is different from the univariate Pearson correlation model, since the latter is computed between each rsFC and actual age, while the former is between each FC and predicted age that involves all the features and is no longer a univariate approach.

In order to understand the impact of statistical model choice (univariate, multivariable prediction with feature filter, multivariable prediction without inversion, multivariable prediction with inversion) on the spatial patterns of rsFC associations with age, we plotted the results from our statistical models on the connectome. Further, we made heat maps to illustrate the percentage of the strong predictive features surviving 1000 random samplings on both days. To be more specific, for each model, we z-scored all the features across all 1000 random samplings on the two days, and thresholded the results by |Z|>2 to create a binarized feature map. Then, for each ROI pair, we counted the number of the sampling rounds (2000 matrices in total for two scan days) that was marked as “1” in the binarization (i.e. |Z|>2) and divided it by 2000 to achieve the survival percentage of this ROI pair as we showed in the heat map.

#### 2.4.5 Test-Retest Reliability

We assessed the reliability of our ML model by evaluating its performance consistency across different subjects and days. Given the samples from two scan days, we first conducted the random sampling and cross-validation on two days separately to obtain two training sets and two test sets. We took the average of the predicted test set on Rest 1 using the trained model from Rest 2 and the predicted ages from the test set on Rest 1 using the trained model from Rest 2 (Figure 2). Since we proposed to compare two different random sampling implementations and two LSVR models with or without feature filters, we would expect in total 12 sets of predicted labels for all participants (two sampling approaches, two feature selection approaches, and two scan days plus one test-retest between scan days).

Intraclass correlation coefficient (ICC) is another standard metric to assess reliability. An ICC statistic centers and rescales observations via a pooled sample mean and standard deviation. Various forms of ICC scores have been carried out by existing literature, and ICC(2,1) is utilized in our study to evaluate the consistency of predicted outcomes from LSVR models for all subjects across two scan days, which is based on the guidelines for ICC scores in the previous studies (Koo & Li, 2016; Taxali et al., 2021). To be more specific, we predicted the labels in the test set on Rest 2 via the LSVR model trained on Rest 1, repeated the same procedures on the training set from Rest 2 and the test set from Rest 1, and then calculated the ICC(2,1) statistic according to the rigorous formula available in the supplementary materials.

### 2.5 Network Level Enrichment Analysis

Network Level Analysis (NLA) was employed to determine patterns of ML results which clustered within specific brain networks. NLA has been previously used to determine brain networks associated with behavioral outcomes by modeling univariate rsFC correlations (Eggebrecht et al. 2017; Marrus et al. 2018; McKinnon et al. 2019; Wheelock et al. 2018; 2019; 2021; 2023). In a novel application, to facilitate accessible biological interpretation of the beta estimates and predictions, we applied NLA toolbox to the results from our ML models which used the features across the entire connectome.

#### 2.5.1 Observed Network-Level Enrichment

We generated four sets of inputs for NLA to evaluate the significant network blocks predictive of age. For the Pearson correlation model, we calculated the mean FC-age correlations across 1000 resampling rounds as the input. For the LSVR model, we used the mean estimated FC-age beta weights across 1000 iterations to be the input. Similarly, for the LSVR model with Pearson feature filter applied ahead, we utilized the mean estimated beta weights across 1000 rounds for the selected features only as the NLA input. Finally, for the LSVR model with the inversion applied, we estimated the inverse weights in each iteration and computed the average across 1000 replications.

For brevity, we refer the four sets of NLA inputs introduced above to edge values in the following description. These edge values were first z-scored, thresholded at |Z|>2 and then binarized (Figure 3A). We selected the threshold value of 2 after experimenting with several thresholds. Among them, 2 provided a balanced distribution of features throughout the connectome, striking a suitable balance between sparsity and density. This particular threshold has been qualitatively evaluated in the previous study (Wheelock et al. 2023) using a significance level of p<0.05 in Pearson correlation analysis. Edges within each network block passing this threshold were used to compute the χ^2^ value relative to the distribution of all edges passing threshold in the rest of the connectome. The 1-degree-of-freedom χ^2^ test was used to compare the observed number of strong (thresholded and binarized) brain-behavior edge values within a pair of functional networks to the number that would be expected if strong brain-behavior edge values were uniformly distributed across the full connectome (Eggebrecht et al. 2017; Wheelock et al. 2021). A large resulting test statistic can indicate that the number of strong associations within a specific network block is enriched, meaning the number of strong edge values is much greater than expected.

**Figure 3.**
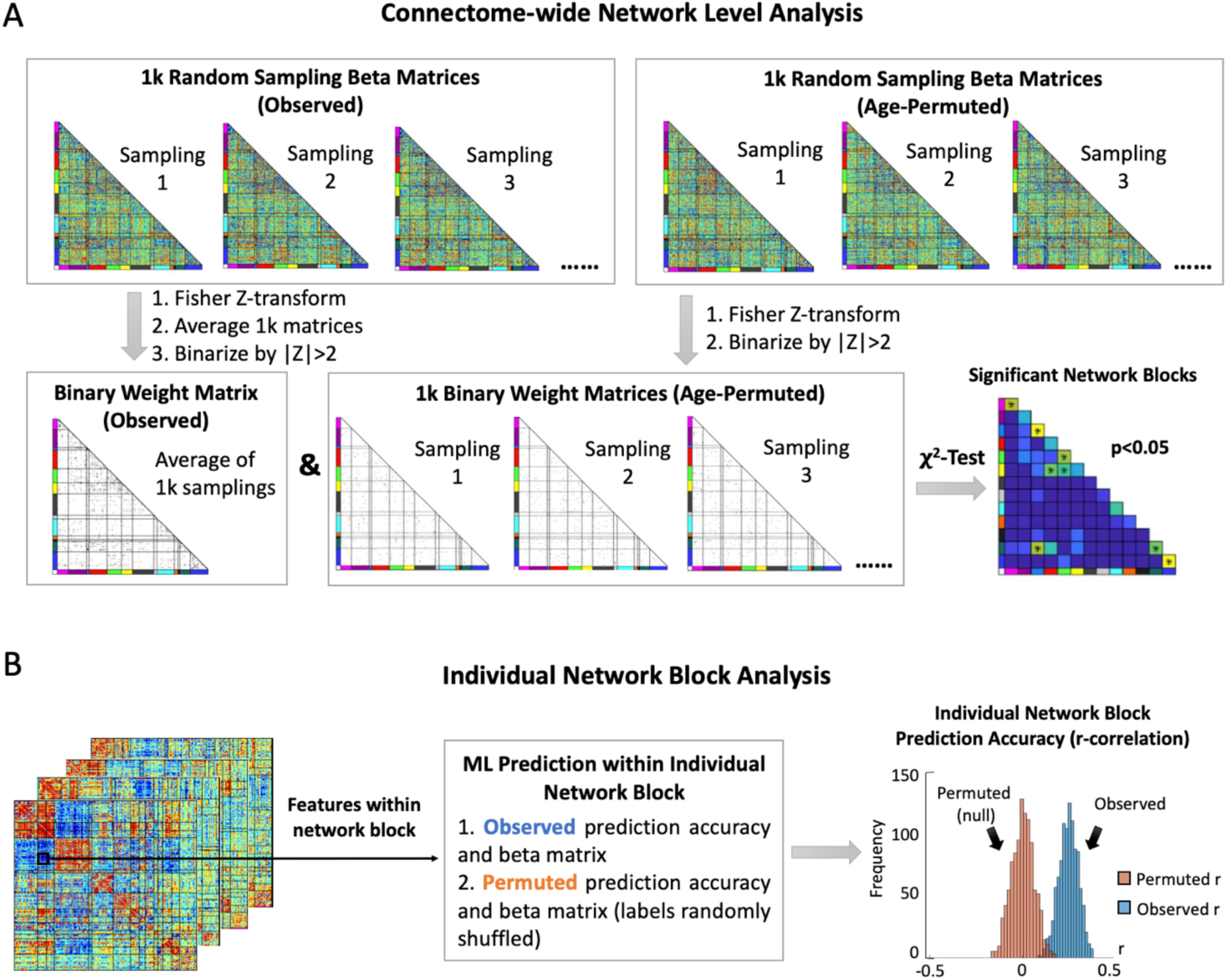
Localizing predictions to biological systems. For the purposes of determining significant network blocks towards prediction and biological interpretation, two different network-block analysis methods were employed as comparison. **(A)** Connectome-wide Network Level Analysis (NLA). The NLA software used the output from the method illustrated in Figure 2 which was repeated 1000 times with response labels (age) randomly shuffled to generate permuted beta weights. Both observed and permuted beta weights were z-transformed and binarized at a threshold of |Z| > 2. These thresholded and binarized Z-values were then used as inputs to a χ^2^ test, yielding a χ^2^ test statistic and p-value. Once statistics and p-values were computed for observed and permuted data, all the p-values from χ^2^ test were concatenated together and then ranked from smallest to largest. The rank of the observed p-value, number of permutations, and number of network pairs were used to calculate an experiment wide permutation-based, network level p-value. The network blocks with this p-value < 0.05 were considered as significant ones towards prediction of age. **(B)** Individual Network Block Analysis. In contrast, for each individual network block, we fit a ML model with the features from within this certain network block by applying the same random sampling and cross-validation implementation as the ML pipeline in Figure 2. We then compared this observed distribution of ML beta weights to a permuted null distribution of weights generated by fitting a ML model with features only from this network block but on randomized age labels. We evaluated the prediction accuracy of each network-block prediction model with the prediction accuracy quantified by correlating the observed and predicted age labels. The same procedures were performed another 1000 times for each model to obtain permuted r-correlations.

#### 2.5.2 Permutation Test

To evaluate the network-level significance of each network block, we adopted the permutation test to compute the permutation-based p-values. To generate the permuted data, we shuffled the age labels and fit the same model (Pearson correlation, LSVR no filter, LSVR with Pearson filter, or inverse LSVR) with the connectome data to create null brain-age weight matrices. This procedure was repeated for 1000 iterations for each model, respectively. χ^2^ test statistics were also calculated on permuted models, generating a null distribution of network-level statistics. The observed (i.e., real) χ^2^ values were then compared to the null distribution to establish network-level significance for the permutation-based p-value < 0.05. Specifically, for each network pair, we computed the permutation-based p-value by ranking the observed network r-correlation between the predicted and actual labels compared to all permuted r-correlations in the entire connectome (lower rank = observed r-correlation is less than more of the permuted r-correlation), and p=1+rank/(1+1000*91), where 1000 is the number of permutations and 91 is the total number of network blocks.

Furthermore, we quantified network-level reliability using Matthew’s Correlation Coefficient (MCC). We first determined the confusion matrix by treating Rest 1 and Rest 2 as observation set and prediction set respectively. The significant blocks in Rest 1 were denoted as “true” and other blocks as “false”. Similarly, significant blocks in Rest 2 were denoted as “positive” and other blocks as “negative”. Accordingly, we can obtain the true positive (TP), true negative (TN), false positive (FP) and false negative (FN). One shall note that denoting Rest 2 as “true” will produce the same result due to the symmetry of MCC scores. With this configuration, we can calculate MCC scores by the formula 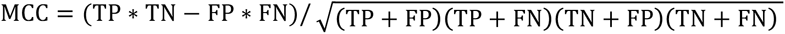. The MCC ranges from 0 to 1 with 1 being more consistent across two scan days.

It is worth noting that many investigators have studied the proper ways to draw a null distribution. See, for example, Xia et al. (2018) shuffled the networks in addition to permuting the labels to generate the null distribution; Zamani Esfahlani et al. (2020) summarized different ways to create a null model that could preserve neuroanatomical realism; and more recently, Váša et al. (2021) provided a comprehensive review on the logic, implementation and interpretation of null models for functional connectomes. Here, we have not shuffled the functional networks of the brain in order to preserve the underlying covariance structure of real biological data (Nichols & Hayasaka, 2016). We examined the impact of shuffling FC networks for the permutation test as in Xia et al. (2018) and observed that the resulting permuted data did not resemble the covariance structure of real data (Figure S2 in Supplementary Materials).

#### 2.5.3 Network-Block Feature Analysis

In contrast to the NLA analysis of connectome-wide ML beta weights applied in this paper, prior work seeking to localize predictive features to biological systems has used only the features within an individual network block for prediction (Millar et al., 2020, 2022; Nielsen et al., 2019; Rudolph et al., 2018). Specifically, different from the NLA which uses all regression weights in the entire connectome from an ML model estimated one time as input, a network-block feature analysis method performed on the within and between-network rsFC of 13 functional networks would fit 78 individual ML models (i.e., an ML model for a network block). For brevity, we call the former NLA method a “connectome-wide” analysis and the latter an “individual network-block” analysis. Specifically, an “individual network-block” test uses the permuted and observed weight matrices from the same network block individually and then compares the distributions of r-correlations between predicted and true labels. If a significant shift is observed between these two r-distributions, then it can be inferred that this specific network block contributes significantly to the prediction of labels (Figure 3B).

While generating a null distribution using permuted labels is one method to determine the significance of a network block, a more stringent evaluation of the biological specificity of a given network block towards prediction could be achieved by comparing to randomly selected features outside of that network block. Given that prediction accuracy is a function of feature set size (Domingos, 2012; Guyon & Elisseeff, 2003; Nielsen et al., 2019), it is important to compare to an equally large feature set selected from the rest of the connectome external to the network block being tested. To evaluate the performance of this “individual network-block” compared to random feature set type network block feature method, we first conducted the “individual network-block” analysis, and for each significant block yielded by this approach, we further generated a different null model which randomly selected the same number of features from the full connectome excluding all the significant blocks. We compared the distributions of r-correlations between the predicted and actual labels by this new null model and the observed block model. If the random model outperforms the observed one, then we can conclude that the “individual network-block” type methods might not be valid for the network block feature analysis.

## 3. Results

### 3.1 Shared variance among families and use of feature filter impacted prediction performance

When comparing the model performances with the random sampling schemes considering the family structure (RS2) and not considering the family structures (RS1), we observed that training the ML model without taking the shared variance among related subjects into account led to the falsely inflated prediction accuracy (r), explained variance (R^2^) and reliability (ICC) (Figure 4A-C). Further, when a marginal Pearson filter was applied ahead of LSVR fitting, we noticed a significant decrease (p-value<0.001) in the prediction accuracy, variance explained, and reliability (Figure 4A-C).

**Figure 4.**
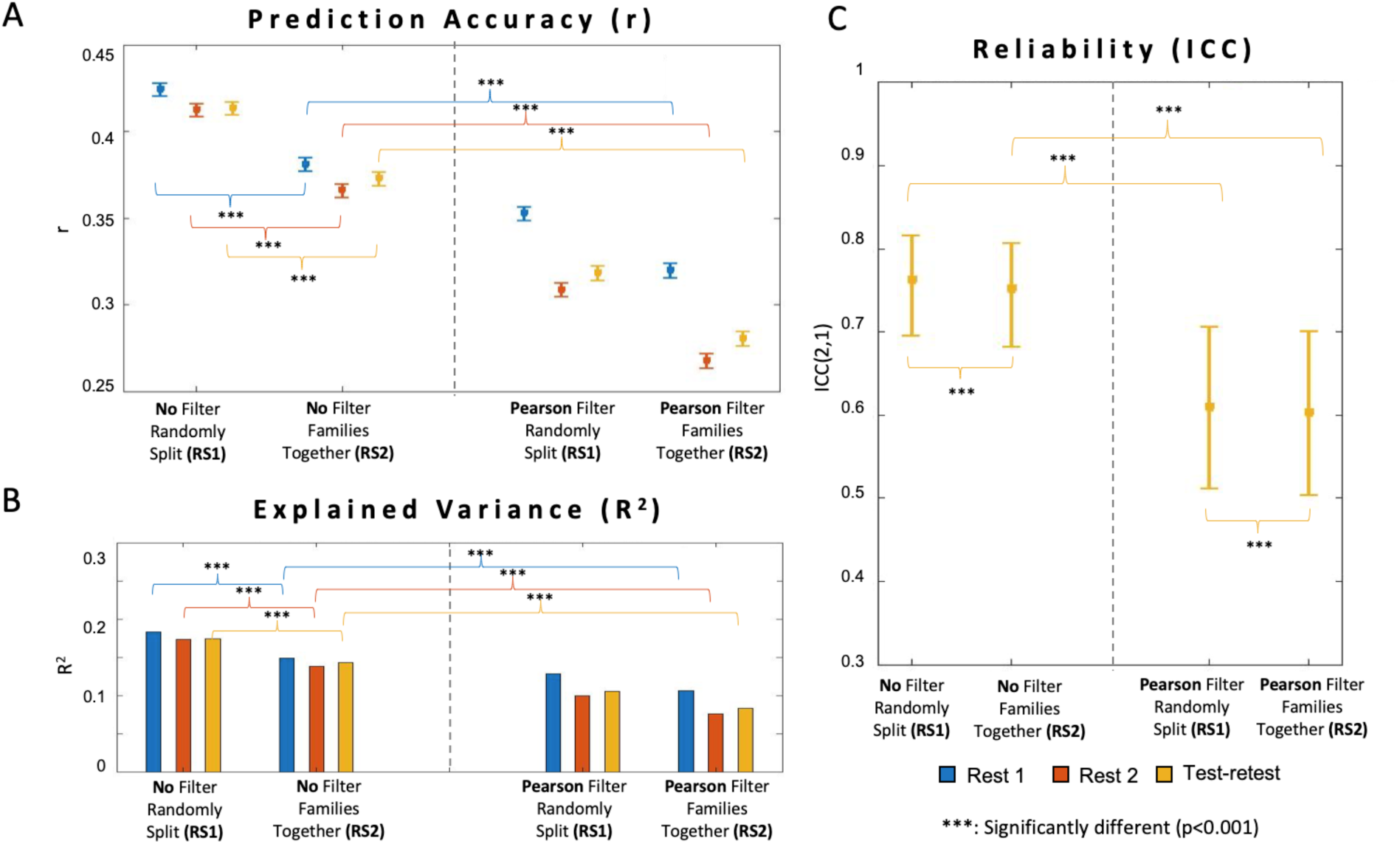
Evaluations of the ML models with two cross-validation sampling strategies and two edge-selection approaches. **(A)** Pearson-r correlation between the actual and predicted labels, with error bars indicating the standard deviation of r correlations. **(B)** The explained variance by the model. **(C)** The reliability of the model quantified by ICC(2,1) for two different days, with error bars indicating the standard deviation of ICC(2,1) scores.

### 3.2 Feature selection via the marginal Pearson filter changed the distribution of ROI with the strongest beta weights from the LSVR model

We investigated the relations of different ROI pairs to the response label (age) via three different measures, including 1) the univariate marginal Pearson correlation between each rsFC feature and age, 2) using the top 1000 ranked univariate Pearson correlations as a feature selection step for LSVR, and 3) all rsFC entered as features predicting age, and 4) all rsFC entered as features predicting age with inversion of the LSVR beta weights. Patterns of strong univariate marginal Pearson correlations between rsFC and age qualitatively appeared to cluster within network blocks along the diagonal and in off-diagonal network blocks including visual-auditory (VIS-AUD) and visual-cingulo-opercular (VIS-CO) (Figure 5A). As expected, we observed that the distribution of features within the connectome with the strongest beta weights obtained by Pearson filter + LSVR appeared in very similar network blocks as the strong univariate Pearson correlations, though, by definition, these results were sparser (Figure 5B). In contrast, the LSVR model on the full connectome (without feature filter) yielded a very different pattern of results across the connectome. Specifically, strong beta weights tended to be clustered in network blocks such as the default mode network (DMN), dorsal attention network (DAN), and frontoparietal network (FPN) (Figure 4C). This showed that a pre-step of feature selection before applying the LSVR model could dominate the estimation results from the ML algorithm. However, strong ML regression weights cannot be interpreted as corresponding to neural predictors of age (Chen et al., 2022; Haufe et al., 2014). Therefore, we applied the inversion to the LSVR model to obtain the multivariable weights in which the directionality and magnitude can be interpreted. We observed that the spatial pattern of rsFC features most predictive of age after inversion were in similar network blocks to those from the univariate marginal Pearson correlation (Figure 4D).

**Figure 5.**
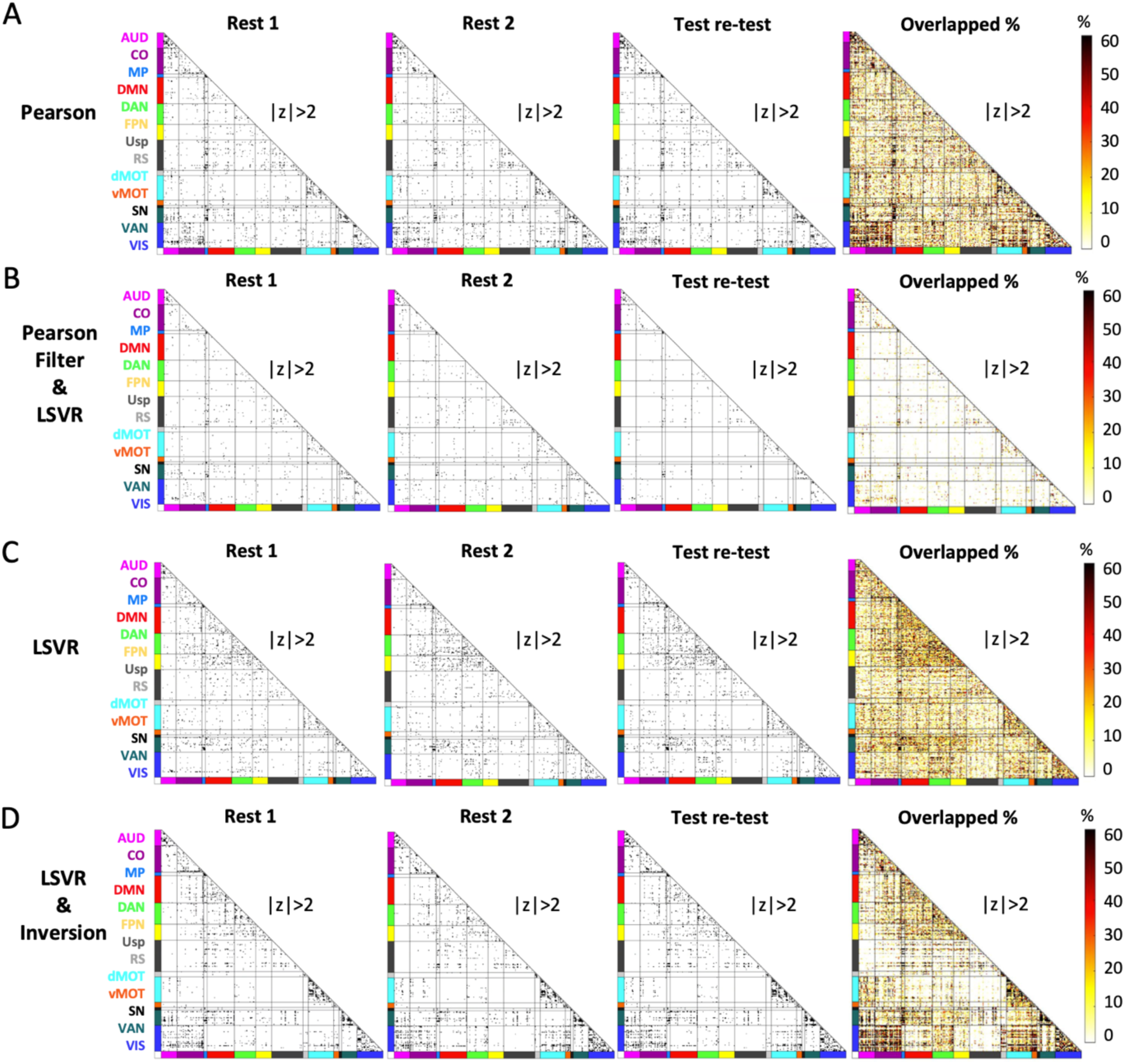
Nominally significant ROI pairs selected by four different methods on two days. **(A)** Pearson-r correlation. **(B)** Pearson-r correlation feature filter and LSVR. **(C)** LSVR with no feature filter. **(D)** Inversion of LSVR model weights without feature filter All results were z-scored, absolute valued, and thresholded at |Z|>2 to standardize analysis and display of rsFC-age associations across analysis methods. Test-retest was calculated by training and testing across two different days. Overlapped % of each ROI over all the random sampling rounds and two scan days were calculated and visualized as heat maps in the last columns of **(A)**, **(B)**, **(C)**, and **(D)**.

### 3.3 The inherent covariance structure of rsFC data should not be removed during the permutation test

For the univariate Pearson correlation, the LSVR models with and without inversion applied, we visually inspected our permuted weight matrices prior to analysis. We observed that randomization of the age labels resulted in permuted Pearson correlations results that were relatively stronger in some network blocks than others, but that the network blocks with these strong associations changed across permutations. In contrast, we observed consistent patterns of strong estimated LSVR weights and inversed LSVR results within consistent network blocks across permutations even with the labels randomly shuffled (Supplemental Figure S2). However, when we randomized the age labels and shuffled the networks in the connectome using the method described in Xia et al. (2018) the resulting permuted connectivity matrices for both Pearson and LSVR models exhibited a relatively uniform distribution across networks and did not retain the underlying covariance structure of the data. Given the lack of biological realism of these weight matrices, the method of randomly shuffling networks was not used for permutation testing in subsequent analyses.

### 3.4 Network level enrichment characterized networks most predictive of age

NLA with four different sets of inputs (i.e., Pearson, Pearson filter + LSVR, LSVR, and inverted LSVR) characterized different network blocks that were most predictive of the label, age. Pearson correlations and the beta weights from the LSVR model with Pearson filter resulted in similar significant network blocks, consisting largely of within-network blocks on the diagonals (Figure 6A, B). In contrast, the NLA with beta weights from LSVR model as input selected distinctly different networks, such as DAN, FPN-DMN, and FPN-DAN (Figure 6C). When applying the inversion technique to the LSVR model for biological interpretation (Chen et al., 2022; Haufe et al., 2014), we observed that it yielded several significant network blocks that were consistent with the marginal Pearson and Pearson filter + LSVR methods and inconsistent with the spatial pattern of betas from LSVR without inversion, including auditory-visual (AUD-VIS) and cingulo-opercular-visual (CO-VIS) (Figure 6D). For each of the four methods, we assessed the network-level reliability by the MCC scores, which were 0.761, 0.640, 0.807, and 0.643, respectively, as shown in Figure 6. The chord plots of the significant network blocks tested by NLA with these four different sets of outputs are presented in Figure 7.

**Figure 6.**
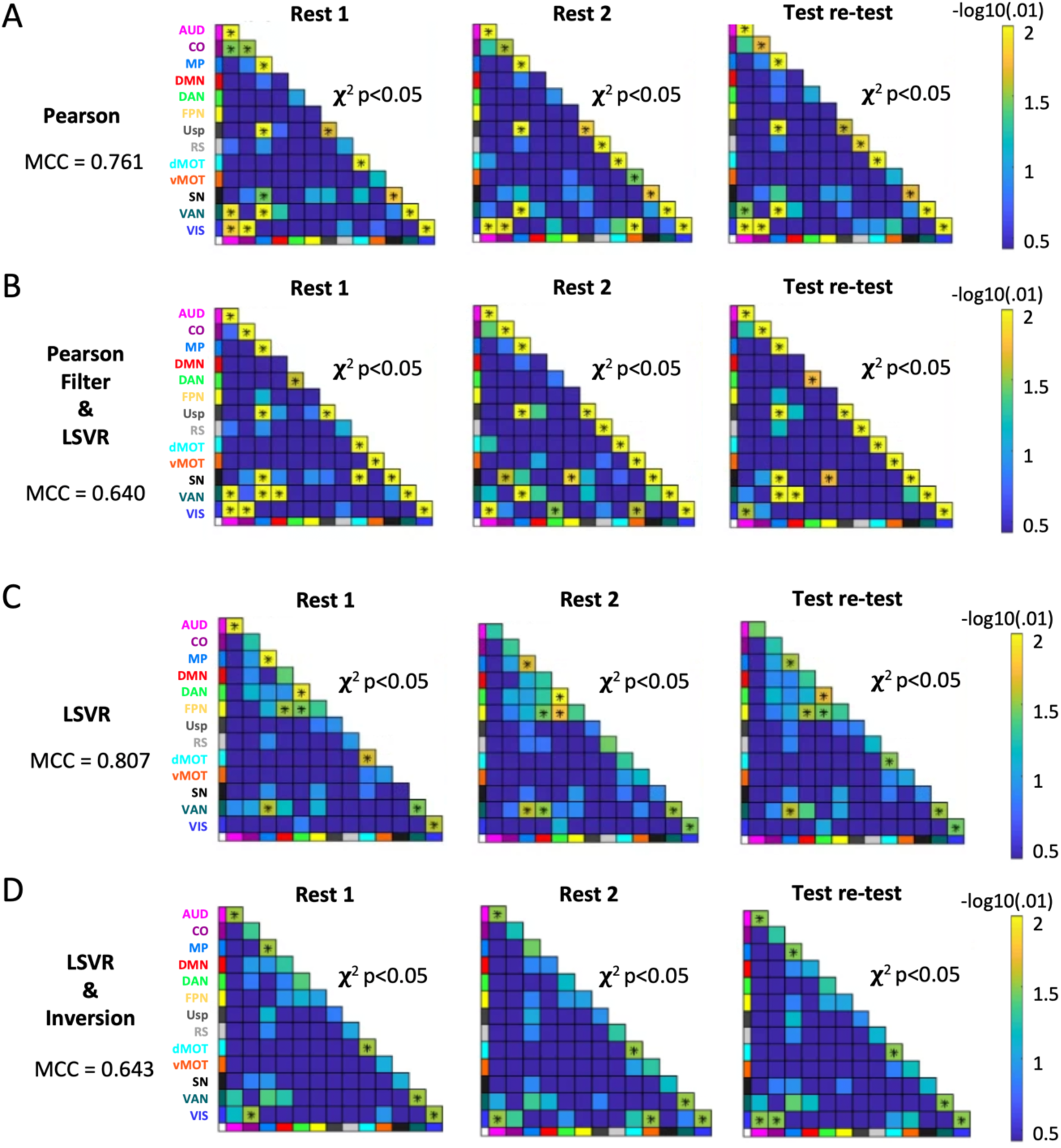
Significant network blocks selected on two scan days by network-level analysis (NLA) with four different sets of inputs. Specifically, the inputs were obtained from **(A)** Pearson r-correlation, **(B)** Linear support vector regression (LSVR) model with Pearson feature selection applied ahead, **(C)** LSVR model, and **(D)** LSVR model with inversion. The Mathews correlation coefficients (MCC) were also calculated for each method.

**Figure 7.**
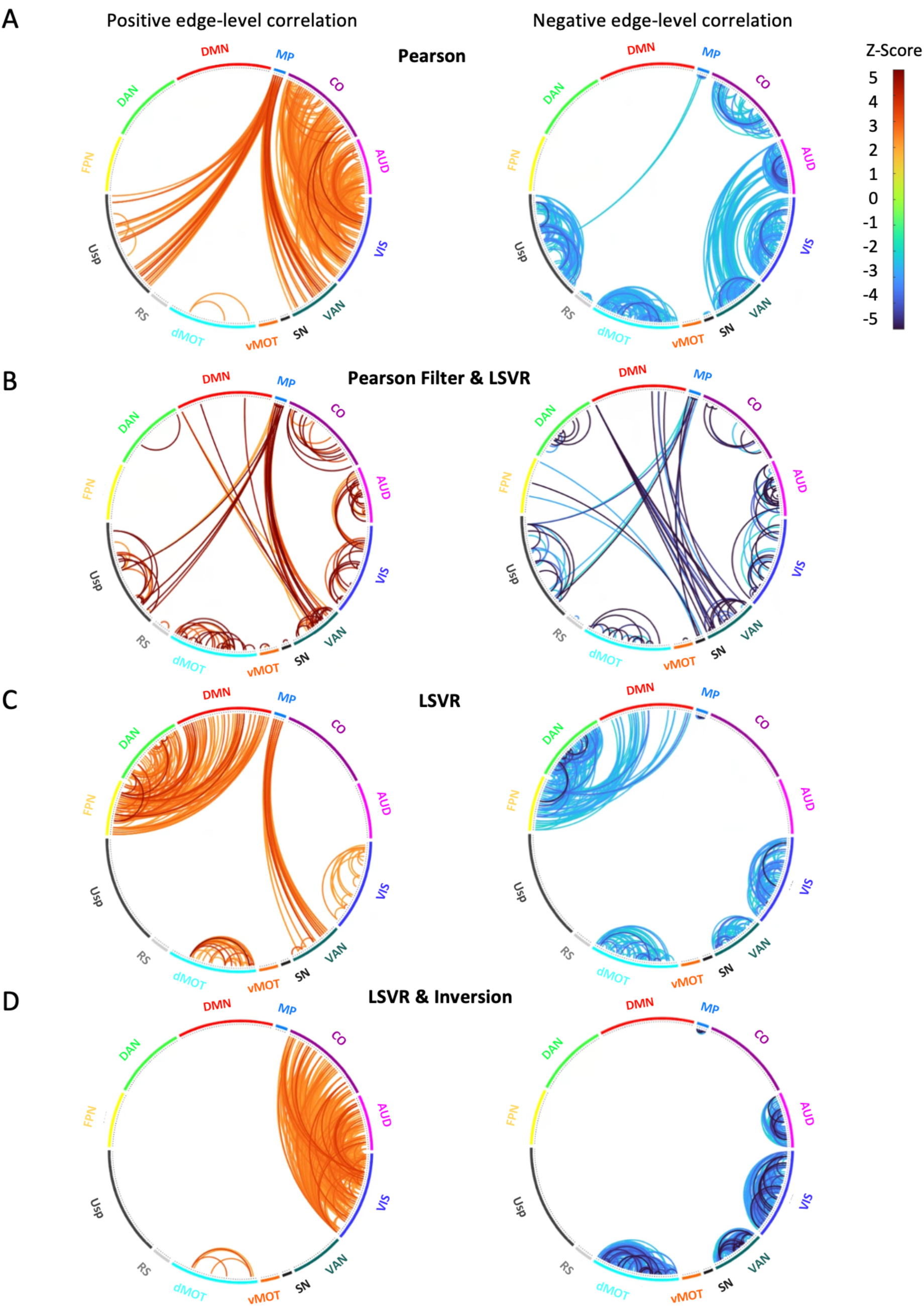
Edge-level chord plots of the significant network blocks selected by the four models. Positive (left) and negative (right) edge-level correlation (p<0.05) are shown respectively within the significant network blocks by the following four models: **(A)** Pearson-r correlation, **(B)** LSVR model with Pearson feature selection applied ahead, **(C)** LSVR model, and **(D)** LSVR model with inversion. All the chord plots correspond to the test-retest column in Figure 6.

### 3.5 Interpretations of significant network blocks cannot totally rely on the network-level ML models

In addition to NLA, we also performed the more standard “individual network-block” analysis which only uses features from within a single network block for each model and has been employed in previous studies for biological interpretation (Nielsen et al., 2019; Rudolph et al., 2018). We observed that almost all the network blocks were significantly predictive of age when compared to a null distribution generated by shuffling the age labels using either permutation-based p-values or Cohen’s D (Figure 8A). While most network blocks were highly significant using this method, we selected a subset of network blocks based on the significant SVR results from the NLA model for illustration purposes. Figure 8B illustrates the large shifts in the distributions of the Pearson r-correlation between actual and predicted ages when we compared the observed and permuted models. While the difference in Pearson r-correlations between actual and predicted ages appeared highly significant, when we instead compared the actual r-correlations within a network block to actual r-correlations from an equal count of randomly selected features outside the network block, we observed that there were no network blocks that predicted age better than randomly selected features (Figure 8A). For illustration purposes, we again selected the same subset of seven network blocks and demonstrate the overlap between network-block prediction accuracy (violin) and prediction accuracy from the same number of features randomly selected outside that network block from the rest of the connectome (scatter) (Figure 8C).

**Figure 8.**
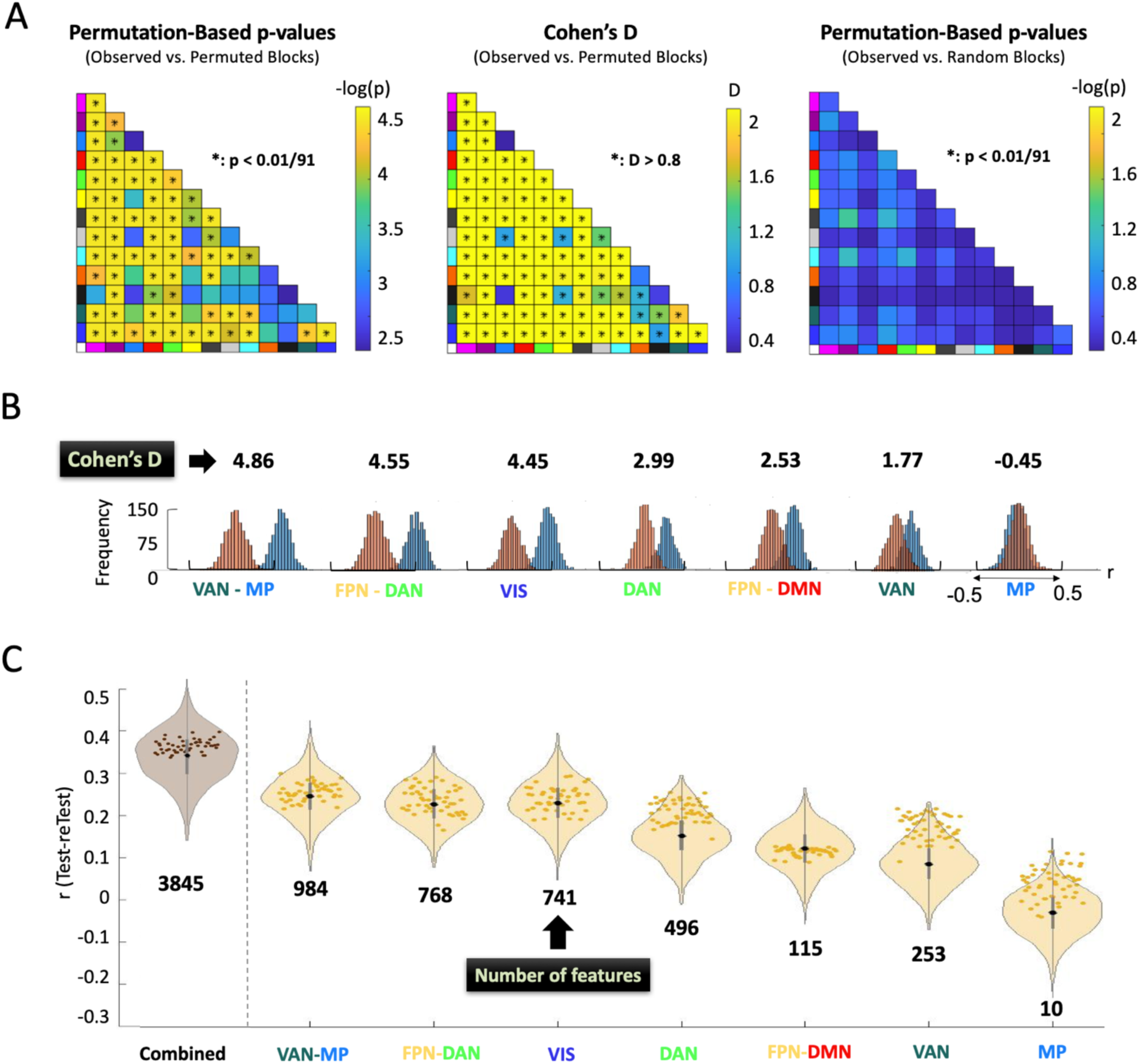
Results of the individual network-block analysis. **(A)** Results of three different hypothesis tests from the “individual network-block” analysis. The Permutation based p values and Cohen’s D values were computed with reference to the null distribution from permuting the age labels. The last plot computed the permutation-based p-values with reference to randomly selected features of the same count without shuffling the age labels. (B) For comparison and illustration purposes, the shifts in the r-distributions of observed (orange) and permuted (blue) models fit with only the features within the individual network blocks (as identified as significant in Figure 6C) are plotted with the corresponding Cohen’s D values measuring the shift between the observed and permuted distributions. (C) For illustration purposes, these same seven network blocks are compared between prediction accuracy of the individual network block ML models (violin) and the prediction accuracy from ML models based on randomly selected features of the same count as the individual network block (scatter). The “combined” violin plot is the ML model fit with the features from all the seven significant network blocks yielded by the “connectome-wide” NLA.

## 4. Discussion

The main goals of this study were to determine the practical application and interpretation of ML models applied to human neuroimaging data. Specifically, we aim to 1) understand the impact of shared variance from related individuals, 2) determine the effect of feature filter step, and 3) apply a novel network level analysis software to increase reliability and biological interpretation of ML results. To this end, we employed LSVR models to investigate the relationship between functional connectome data and age. First, we tested a random resampling procedure to control shared variance among related subjects and observed that ignoring the shared variance resulted in falsely inflated prediction accuracy. Further, we compared the ML models with and without feature selection applied ahead of model fitting and found that a marginal Pearson feature filter resulted in lower prediction accuracy and test-retest reliability. Finally, we demonstrated that an accessible biological interpretation of ML models can be provided by utilizing the NLA method and ML model inversion while ML models fit within a single network block were not better than randomly selected features at predicting age.

### 4.1 Separate related individuals into training or test sets but do not allow in both

A rising number of researchers are applying ML models to predict behavioral or clinical outcomes using the functional connectome. However, few prior studies have examined standardized methods for modeling shared variance among families (Cohen et al. 2020; Elliott et al. 2019; Feilong et al. 2021). One issue introduced by cross-validation procedures in datasets with related individuals is that the related subjects could appear in both the training set and hold-out test set. Given that related individuals and twins have connectome data that are more similar than between unrelated strangers (Demeter et al., 2020; Miranda-Dominguez et al., 2018), the shared variance among families violates the assumption that cross-validation is performed on an independent dataset. As a direct consequence, when predicting the labels in the test set, since there are related subjects from the training set whose labels are known, the prediction accuracy yielded by the test set cannot be used as a reliable measure for the model performance. In other words, this type of cross-validation fails to avoid overfitting of the ML models. Therefore, based on the cross-validation method proposed by Feilong et al. 2021, we developed a random sampling regime considering family structure to avoid this drawback. While prediction accuracy and reliability were lower when keeping the subjects from the same family together within either training or test set, we can better guarantee that the prediction in the hold-out test set is not contaminated by the information in the training set, and therefore prevented the falsely inflated prediction accuracy. Our results have implications for existing datasets with family members such as HCP, ABCD, IBIS, Dominantly Inherited Alzheimer Network (DIAN), etc. and individuals who analyze these datasets should use a cross-validation approach keeping families separated to either train or test datasets.

### 4.2 Univariate feature filters can lead to low prediction accuracy and reliability in the presence of the correlation among regressors

Various feature selection methods have been applied ahead of the regression step due to the computational cost of high-dimensional data (Dadi et al., 2019; Nielsen et al., 2019; Shi et al., 2021). Among these methods, the marginal Pearson correlation is a popular option in functional connectome research (Nielsen et al. 2019; Fan et al. 2006; Shen et al. 2017). However, based on our study, we found that this additional screening step reduces model performance. Specifically, reducing features via marginal screening methods results in lower prediction accuracy and reliability and different networks associated with outcome prediction or, in the current study, age. This is not simply due to the decrease in the number of features entering the model since the accuracy of prediction tends to converge when the number of features exceeds 1000 (Nielsen et al., 2019). The suboptimal performance of univariate feature selection methods, such as marginal Pearson correlation, can be attributed to their tendency to select a multitude of strong edges that may arise from redundant correlations among features. For instance, if the connectivity between two networks is correlated with age, the Pearson filter will tend to pull many features within that network block. Hence, the information about the relationship between rsFC and age is largely redundant across features within a network block (i.e., all rsFC tend to be positive or all negatively correlated with age to varying degrees). In contrast, multivariable approaches, like utilizing inversed feature weights from a predictive model such as LSVR, have the advantage of selecting unique and impactful rsFC features that contribute significantly to the prediction, irrespective of correlations among features. Consequently, when relying solely on univariate correlations to select features, those features that could have been crucial for prediction are likely to be eliminated during marginal screenings (Wang et al. 2020). In our study, we observed that the significant network blocks chosen by Pearson Filter + LSVR appeared to be more similar to the ones by simple Pearson correlation than they did to the ones by LSVR models. The Pearson filter tends to select many redundant features that are correlated among themselves but not necessarily significant for prediction to enter the LSVR model. Also, the redundant features for the ML model are often located within the network blocks. These redundant features would have been removed during the regularization with L^2^-penalty in the full LSVR model. Alternative feature selection methods that account for multivariable relationships between features such as the iterated sure independent screening (ISIS) (Fan and Lv 2008), and the correlation measures including tilted correlation (Cho & Fryzlewicz, 2012) and quantile partial correlation (Ma et al. 2017) may have greater utility in human connectome analysis.

### 4.3 LSVR and NLA on the inverted weights rather than the regression weights provide an accurate and biologically interpretable framework for predictive ML models

The performances of LSVR models without feature filters were consistent with the existing age prediction literature in both prediction accuracy and reliability (Cui & Gong, 2018; Taxali et al., 2021). Even after correcting the falsely inflated effect caused by the shared variance, our ML model still had competitive prediction accuracy and reliability. We further applied NLA software to determine which network blocks were more associated with age than the rest of the connectome. NLA methods were previously developed for univariate FC association studies (Eggebrecht et al. 2017; Wheelock et al. 2018; 2019; 2021). Here for the first time, we demonstrate their utility in a multivariate setting. Regarding the contribution to the prediction of age, NLA revealed that LSVR coefficients were consistently (across days by test-retest model) stronger and more clustered within several networks including medial parietal (MP), ventral attention (VAN), dorsal attention (DAN), and frontoparietal (FPN) relative to the rest of brain. Similar to previous literature exploring age associations using linear models and massive univariate analysis (Rieck et al., 2021), we observed a decrease in FC within executive control networks was reduced as age increased, which aligns with our findings presented above. Furthermore, consistent to the prior research focusing on the network blocks that are most predictive of age using ML models (Dosenbach et al., 2010; Rieck et al., 2021), we also identified the right anterior prefrontal cortex as one of the regions with the highest relative prediction power for age. This region is known to play a crucial role in cognitive control and higher-order. In terms of the reliability of our network-level observations, it is noteworthy that the utilization of MCC scores not only enhances the biological interpretability of our findings, but also yields trustworthy estimates of network-level significance.

Furthermore, concerning the reproducibility of NLA results across three different input sets (i.e., Pearson, Pearson + LSVR, LSVR), we observed some similarities such as medial parietal (MP), ventral attention (VAN), visual (VIS) and ventral attention - medial parietal (VAN-MP) that were implicated in all three methods and two scan days as well as the test-retest. However, considering the similarities between the results by Pearson and Pearson + LSVR, LSVR identified a distinctly different pattern of networks. The possible reason is that the LSVR model tends to put more weight on the features with higher deviation across subjects for a better explained variance and higher prediction accuracy, which yields an optimal combination of features for the purpose of prediction (James et al., 2021). In contrast, Pearson and Pearson + LSVR models selected ROI pairs (and by definition the network blocks) with higher individual rsFC relation to the response label given the massively univariate method of analysis.

To achieve a compromise between the two types of interpretation (i.e., univariate correlation and multivariate regression), we applied the inversion model to the ML pipeline by evaluating the covariance between the rsFC and predicted label obtained from the LSVR model. We observed that the inversion model removed the significant network blocks selected by LSVR due to the high variance across subjects, including dorsal attention (DAN), frontal-parietal - default mode (FNP-DMN) and frontal-parietal - dorsal attention (FNP-DAN), while retaining the network blocks selected by univariate type of methods such as auditory-visual (AUD-VIS) and cingulo-opercular - visual (CO-VIS). From the ROI-level heatmaps, it is more explicit that the inversion model yields a similar significance distribution as the Pearson correlation, but with more concentration on VAN, VIS, AUD-VIS and CO-VIS.

We acknowledge that the MCC score for the LSVR model is marginally higher compared to the LSVR model with inversion, despite both exhibiting a strong positive correlation. Nevertheless, it is important to distinguish between prediction and biological interpretation as distinct objectives. The beta weights obtained directly from the predictive LSVR model solely reflect the impact of rsFC on label prediction and do not possess any inherent biological significance. Therefore, we highlight that the inversion model provided a more precise biological interpretation by preserving the individual effects of each feature while also considering the full model since the computation of predicted labels involved all the features (include Haufe citation). This inversion step is crucial for any future research with a goal of biological ML model interpretation rather than prediction accuracy. Alternatively, for the purpose of interpretation, one can consider employing other models that prioritize interpretability, such as the generalized linear model (GLM). Unlike the LSVR models which often rely on multivariable analysis, a GLM is typically referred to as massively univariate in the sense that it is estimating the effect of age or other covariates at each individual FC edge, while a multivariable ML model is fitting age using all the FC edges as regressors simultaneously. We shall clarify the difference between the multivariate and multivariable models. Multivariate models always focus on relationships between multiple dependent variables by allowing multiple response variables simultaneously (e.g., age and other behavioral scores), taking into account the interdependencies and relationships between them. Differently, multivariable models predict a single outcome (e.g., age) via multiple predictor variables. In this study, we mainly compared the multivariable models (e.g., LSVR) to the univariate ones (e.g., Pearson filter). We observed that multivariable approaches allow for the examination of individual variables while considering their relationship with the outcome of interest, thereby expanding the options available for conducting interpretive analyses.

### 4.4 NLA on all features (full connectome) from the ML model is recommended rather than creating individual models containing features from singular network blocks

Besides applying the NLA toolbox to investigate the significance of all the network blocks towards the age prediction, we also utilized the methods introduced in the existing literature (Millar et al., 2020, 2022; Nielsen et al., 2019; Rudolph et al., 2018), where they fit the ML models inside a certain network block and tested the significance of this block by evaluating the shifts between the Pearson r-distribution of the observed and null models. Consistent with prior research (Nielsen et al., 2019), we found that although the distribution of r-correlations between actual and estimated age for each significant network pair were significantly shifted compared to the permuted null, these network-level models performed no better than models created from an equal number of randomly selected features from the rest of the connectome. This suggests that ML models generated on each individual network block cannot be used to infer biological specificity of associations with behavior or clinical outcomes. Similar to prior research, we also demonstrated that the prediction accuracy is a function of feature size where the r-correlation between the predicted and actual labels asymptotically increases with the increasing feature count (Nielsen et al., 2019). However, even when we combined features from multiple network blocks together that appear to be highly predictive in the LSVR NLA model, these >3000 features were not able to perform better than randomly selected features of the same count selected from the rest of the connectome. Consequently, we point out that attempting to model individual network blocks in isolation would fail to provide biological interpretations of the ML results for three reasons: (1) “individual network-block” analysis does not result in prediction accuracy above randomly selecting features as prior work has demonstrated (Nielsen et al., 2019), (2) prediction accuracy is dependent on the size of the network, making different networks incomparable if modeled individually (Millar et al., 2022; Nielsen et al., 2019), and (3) the networks which have greater prediction accuracy than the null are not biologically interpretable because they lack the model inversion and the inversion cannot be applied on an individual net to yield its contribution among all the network blocks (the inversion should be applied on the whole connectome to account for the relationship between different network blocks). We highlight that ML models utilizing whole-connectome data with the hypothesis testing framework of NLA provides a more robust avenue for biological interpretation of ML predictions instead of fitting separate models on the features within each individual network-block.

### 4.5 Shared variance should be modeled with the biological realism maintained in permuted beta weight matrices

In the permutation test conducted for the network-level analysis, we randomly shuffled the age labels across subjects and then estimated the beta weights through these null models. Prior research suggests that permuted models might not be suitable for the null distribution when data do not respect exchangeability under the null hypothesis, such as the tests with known dependence among subjects (e.g., family members) and instead a block permutation would be more appropriate (Winkler et al., 2014). Following these guidelines, we randomly shuffled the age labels across the families, while still keeping the family members together within either the training set or the test set, i.e., viewing each family as a block in our permutation test. We observed that the permuted beta weight matrices generated by LSVR models displayed a consistent structural pattern across multiple sampling iterations. Interestingly, this pattern closely mirrored the standard deviation observed in the functional connectivity matrices across all subjects. Conversely, the permuted brain-behavior matrices produced through Pearson correlation did not exhibit similar structural patterns across permutations. This intriguing observation offers potentially valuable insights into the underlying reasons for LSVR’s successful performance in predictive analysis, and warrants further investigation into how these consistent structural patterns contribute to prediction accuracy.

Furthermore, prior work has suggested that the rsFC features should be permuted in addition to permuting the labels (Xia et al. 2018). We observed that, while a model that shuffles both networks and labels may yield a higher degree of randomness, it risks losing the intrinsic covariance structure of the connectome. Consequently, null models generated in this way might lead to questionable significance results in subsequent hypothesis testing due to potential loss of essential details and information. In the voxel-wise literature, in the presence of spatial dependence among voxels, random sampling cannot be performed independently voxel by voxel, but rather entire images should be resampled as a whole to preserve such dependence structure (Nichols & Hayasaka, 2016). Hence, we recommend future researchers analyzing the connectome only shuffle the behavior or clinical outcome labels and preserve the covariance structure of the connectome for biological realism during permutation testing.

### 4.6 Limitations and future directions

One limitation of this paper is that all analyses were validated on the HCP dataset, and it is crucial to extend our analysis to other datasets to obtain a more comprehensive scope and impact of our findings. In addition, considering the imperfectness of marginal feature filters, alternative feature filter methods should be investigated that incorporate multivariate feature selection. Finally, we only reported LSVR results in this study because we were iterating on so many other methods. However, other models have been used in connectome data that have shown promise, such as Canonical Correlation Analysis (CCA) (Mihalik et al., 2022), deep artificial neural networks (DNN) (He et al., 2020; Niu et al., 2020), random forest (Cohen et al., 2020; Jollans et al., 2019), etc. Additional work is needed to validate NLA with these alternative ML models and determine how these models and corresponding filters impact biological interpretation of prediction weights.

## 5. Conclusion

The present study proposed a sampling scheme to implement ML models accounting for related subjects such as siblings and family members. In the presence of complex correlation structure among the regressors, feature selection approaches based on marginal correlation (e.g., Pearson) could preserve redundant features and rule out highly predictive features thus harming the model performance. Besides the accuracy of predictive models, emphasis on the specific contribution of each brain network cannot be negligible and should be carefully assessed by leveraging ML models with whole connectome data and considering the inversion model for biological interpretations.

## Contributions

Conceptualization: JL, MDW, ATE

Methodology: JL, MDW

Software: JL, MDW, AE

Validation: JL, MDW

Formal Analysis: JL, MDW, AS

Investigation: JL, MDW

Resources: MDW, KK, JCT, BA

Data Curation: MDW, KK, JCT, BA

Writing - Original Draft: JL, MDW

Writing - Review & Editing: JL, MDW, ATE, AS, KK, LC, JCT, BA, NRK, XF

Visualization: JL, MDW, AS

Supervision: MDW

Project Administration: JL, MDW

Funding Acquisition: MDW

## Citation Diversity Statement

Recent work in neuroscience has identified a bias in citation practices such that manuscripts written by women and other minorities are under-cited relative to the number of such papers in the field (Dworkin et al., 2020). Here we quantify the citation diversity of the present manuscript excluding self-citations of the first and last authors of this manuscript. Our reference list contains 52% man-man (first author-last author), 14% woman-man, 22% man-woman, and 12% woman-woman citations.

## Acknowledgements

Study funding support was provided by NIH grants: K99/R00 EB029343

## Data Availability

Human Connectome Project data are available at https://db.humanconnectome.org/.

## Code Availability

https://github.com/WheelockLab/NetworkLevelAnalysisBeta https://github.com/WheelockLab/MachineLearning_NetworkLevelAnalysis

