## Supplementary Materials for "Network level enrichment provides a framework for biological interpretation of machine learning results"

***Linear Regression Model***

For N=965 subjects, we observed the ages $y_1, y_2, \dots, y_n$, and the feature vectors $X_1, X_2, \dots, X_p$, where $X_j = \left(x_{1,j}, x_{2,j}, \dots, x_{N,j}\right)^T$ with $x_{i,j}$ being the observed j-th rsFC for the i-th subject. In each random sampling round, we assigned 80% family to the training set, and fit the linear regression model as follows:

$$\hat{y} = \sum_{j=1}^{p} \hat{\beta}_j X_j + \hat{\beta}_0,$$

where the vector $\hat{y} = \left(\hat{y}_1, \hat{y}_2, \dots, \hat{y}_{n\_train}\right)^T$ consists of predictive labels (i.e., age) of all the family members in the training set, and $\hat{\beta}_j$ is the estimated beta weight for the j-th feature in rsFC. Our goal is to train the above ML regression model and find optimal $\hat{\beta}_j$, $\mathrm{j} = 1, 2, \dots, 55278$, and $\hat{\beta}_0$ such that this model could best predict the actual ages $y_i, i = 1, \dots, 965$. Since the number of features exceeds the sample size, we applied the Linear Support Vector Regression (LSVR) to deal with the high-dimensional problem and estimated these regression coefficients.

***Cross-Validation for Tuning Parameters***

Note that to handle the dimension issue, LSVR model incorporates two parts in the loss function, which are the regularized L2-norm term aggregating all the beta weights and another Vapnik's $\varepsilon$-loss function. We follow the standard construction of these two terms as applied in the previous study (Cui and Gong 2018) and shall save the details here due to the relevance. However, a tuning parameter balancing these two terms is not trivial to derive. In order to find the optimal tuning parameter, we inserted a 5-fold cross-validation in each 1000 random samplings, which means that to implement the LSVR approach for the model fitting, we applied a nested cross-validation method. Specifically, for each outer-sampling round, we randomly assigned 80% family to the training set and the remaining 20% to the test set. For each inner 5-fold cross-validation, we worked on the training set from the outer sampling and continued to split it into a training set and a test set to find an optimal tuning parameter by minimizing the loss function mentioned above. The details are shown in Figure S1.

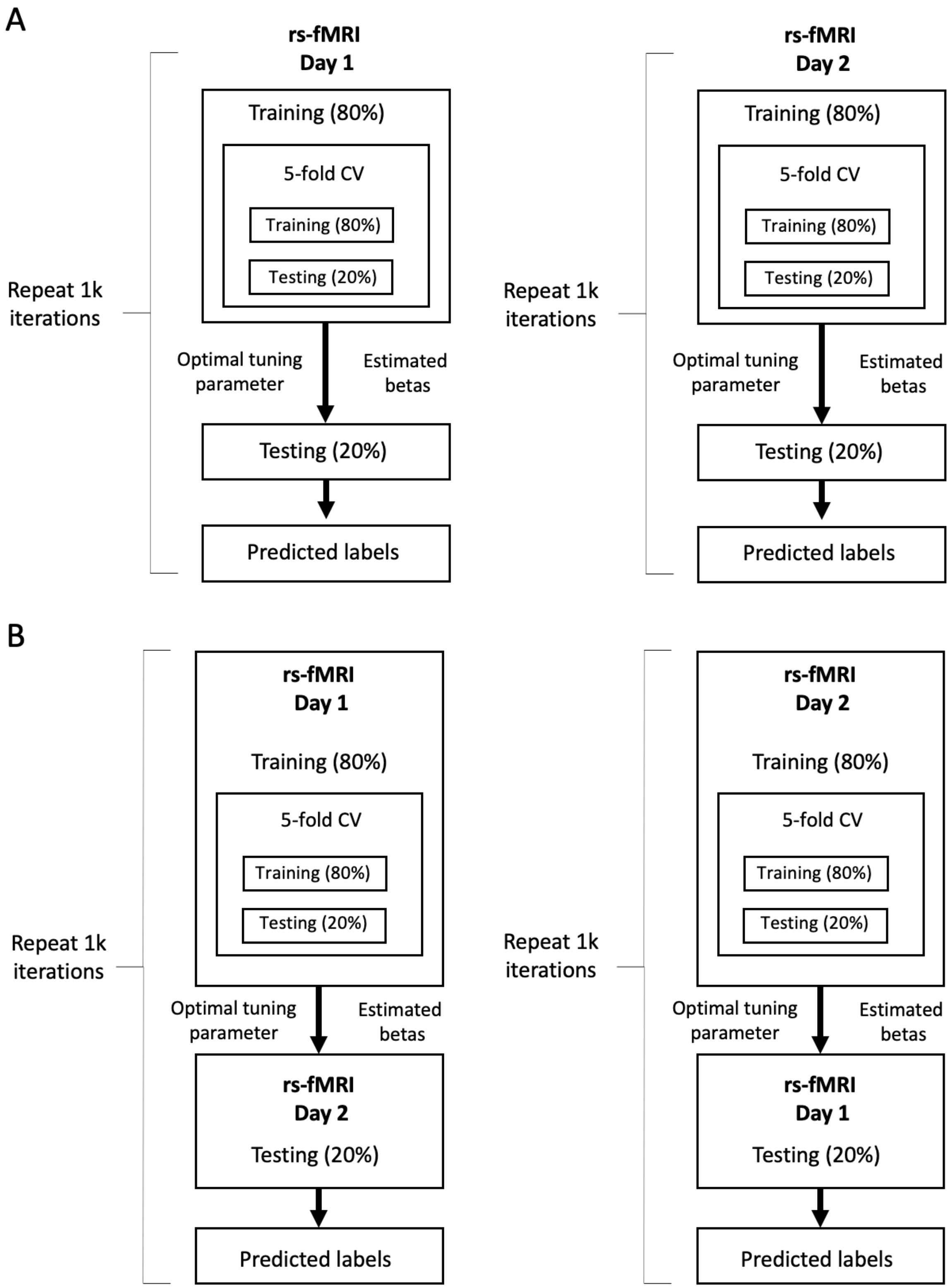


**Figure S1. The nested 5-fold cross-validation (CV) for the ML pipeline. (A)** The nested 5-fold CV on two separate rest scan days. **(B)** The nested 5-fold CV for the test-retest.

***Prediction Accuracy of ML Approaches***

Aside from the correlation between the predicted labels and the actual ones, we provide another two commonly used measures, Mean Absolute Error (MAE) and Mean Square Error (MSE) (He et al. 2020; Niu et al. 2020; Modabbernia et al. 2021) to evaluate the prediction accuracy of four ML approaches with different feature filters applied ahead and different random sampling methods utilized for the regression model fitting. Table S1 showed the MAE and Table S2 presented the MSE, from which we could draw two conclusions: (1) if the shared variance among siblings/families were neglected, then the prediction accuracy of the ML model would be falsely inflated; (2) Margin Pearson filter applied ahead of the ML model could harm the prediction accuracy.

**Table S1** Mean Absolute Error (MAE) of four ML approaches.

| **MAE** | No Filter | | Pearson Filter | |
|---|---|---|---|---|
| | RS1 (Randomly Split) | RS2 (Families Together) | RS1 (Randomly Split) | RS2 (Families Together) |
| Rest 1 | 3.1722 | 3.2391 | 3.1878 | 3.2617 |
| Rest 2 | 3.2221 | 3.2761 | 3.2804 | 3.3560 |
| Test-retest | 3.1970 | 3.2457 | 3.2657 | 3.3266 |

**Table S2** Mean Square Error (MSE) of four ML approaches.

| **MSE** | No Filter | | Pearson Filter | |
|---|---|---|---|---|
| | RS1 (Randomly Split) | RS2 (Families Together) | RS1 (Randomly Split) | RS2 (Families Together) |
| Rest 1 | 15.2517 | 15.8611 | 15.6208 | 16.3052 |
| Rest 2 | 15.6774 | 16.1364 | 16.5109 | 17.2508 |
| Test-retest | 15.4207 | 15.8931 | 16.4068 | 16.9566 |

***Permuted Beta Weight Matrices***

When we performed the permutation test and estimated the beta weight matrices of permuted models with labels (i.e. ages) randomly shuffled, the permuted weight matrices still had the structural patterns. In (Xia et al. 2018), besides the labels, they also permuted the networks to generate the null models. We showed the permuted matrices by these two approaches and compared the patterns. In Figure S2, we list five permuted matrices estimated weight matrices from the ML model with a) labels randomly shuffled, and b) both labels and networks randomly shuffled. For these two implementations, we estimated the beta weights via two methods: 1) calculating the Pearson correlation between each feature and the age; 2) estimating the regression coefficients by utilizing the LSVR model with no feature filter applied. We observed that if we interrupted both the feature and label orders, no structural pattern was captured in the permuted beta matrices. However, when only permuting the labels without shuffling the networks, the structural patterns appeared in each random sampling loop. When the Pearson method is applied, the structural patterns were different from each other, while the patterns from the LSVR models (Column 2 in Figure S2) reflect the underlying variance structure of the rsFC for all subjects. We shall investigate this phenomenon in our future study.

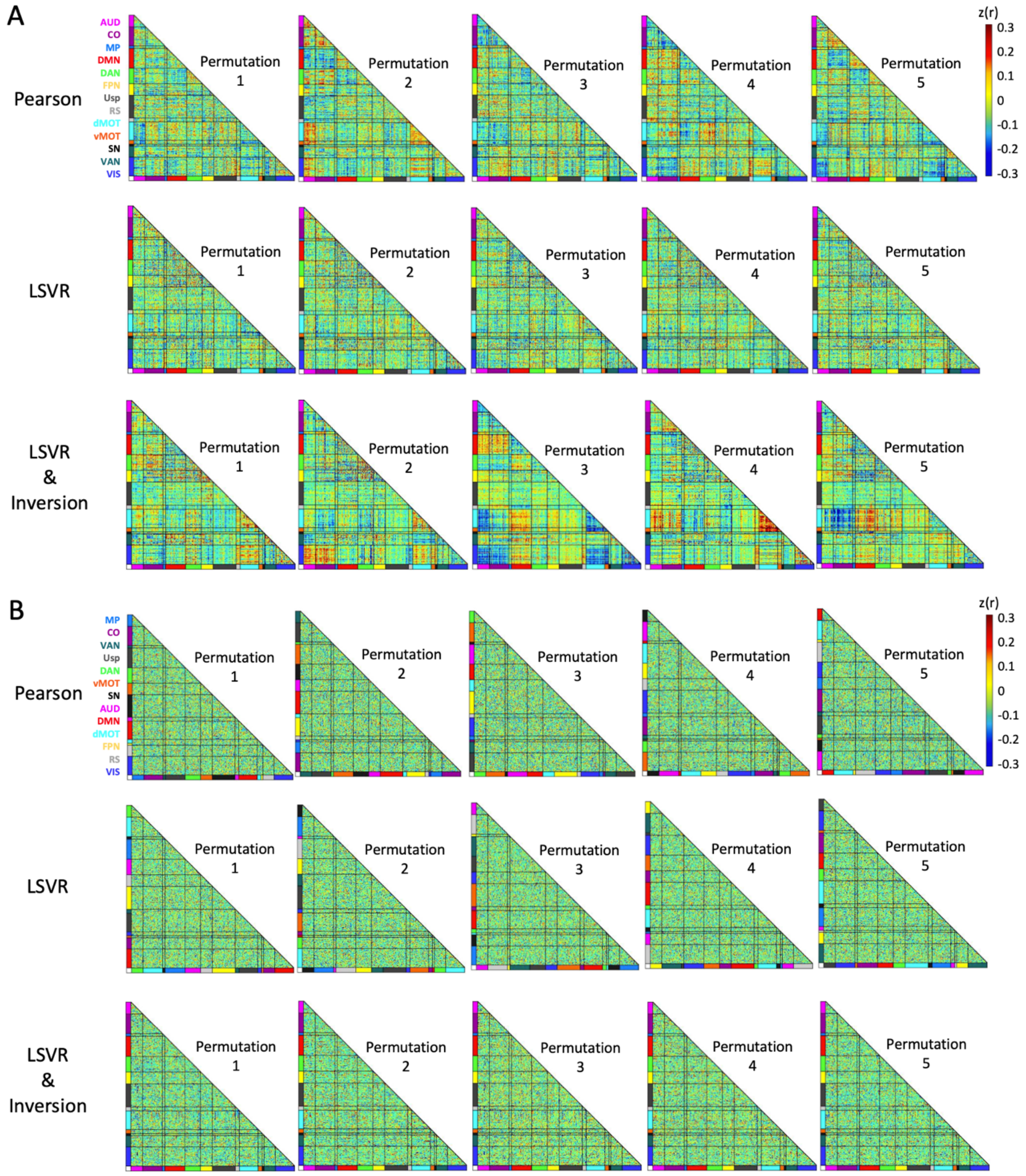


**Figure S2.** The first five permuted estimated weight matrices from the ML model. We examined three estimation methods: Pearson, LSVR and LSVR with inversion. **A.** Only age labels were randomly shuffled. **B.** Both labels and networks were permuted.
